# Learning What a Good Structural Variant Looks Like

**DOI:** 10.1101/2020.05.22.111260

**Authors:** Murad Chowdhury, Ryan M. Layer

## Abstract

Structural variations (SVs) are an important class of genetic mutations, yet SV detectors still suffer from high false-positive rates. In many cases, humans can quickly determine whether a putative SV is real by merely looking at a visualization of the SV’s coverage profile. To that end, we developed Samplot-ML, a convolutional neural network (CNN) trained to genotype genomic deletions using Samplot visualizations that incorporate various forms of evidence such as genome coverage, discordant pairs, and split reads. Using Samplot-ML, we were able to reduce false positives by 47% while keeping 97% of true positives on average across several test samples.

## 1. Introduction

Structural variants (SV), which include large (≥ 50 base pairs) deletions, duplications, insertions, inversions and translocations, are responsible for most variation in the human population and cause a number of genetic diseases (Weischenfeldt et al., 2013). Unfortunately, SV callers often suffer from a high false positive rate, so steps must be taken to filter SV call sets.

Many SVs can be validated by manually inspecting the aligned reads around the called region. Samplot (Layer, 2020) was developed for such a purpose and provides a visualization of alignments in a given locus for a set of samples. For samples sequenced by Illumina paired-end sequencing, Samplot incorporates multiple forms of evidence to help determine the validity of a putative SV call:

1. **Genome coverage** The number of reads aligned to the reference genome at each position across the region of interest. Low signal in the region spanning the reported breakpoints can be evidence of a deletion.
2. **Discordant read pairs** Paired-end reads that deviate too far from the mean insert size. Clusters of these discordant pairs often span an SV breakpoint.
3. **Split Reads** Reads with sequences that map to different regions of the reference genome. These reads also cluster around breakpoints and confer better spatial resolution than discordant pairs.

Identifying a reported deletions as a false positive becomes easy and fast with these visualization tools (Belyeu et al., 2018). Unfortunately, manual curation of SV callsets is simply not feasible since typical SV callsets can contain thousands of regions. In this paper, we present Samplot-ML, a convolutional neural network (CNN) model built on top of Samplot to be able to automatically genotype putative SVs. While Samplot-ML inherently supports any SV type, the current model only includes deletions. There are too few called duplications, insertions, inversions, and translocations in the available data to train a high-quality model. For example, the 1,000 Genomes Project phase 3 SV call set included 40,922 deletions, 6,006 duplications, 162 insertions, 786 inversions, and no translocations. We expect this limitation to be temporary. The workflow for Samplot-ML is simple. Given a whole genome sequenced (WGS) sample (BAM or CRAM) as well as a set of putative deletions (VCF), images of each region are generated using Samplot. Samplot-ML then re-genotypes each call based on its image. The result is a call set where most false postives are flagged. Using Samplot-ML, we demonstrate a 47% reduction in false positives while keeping 97% of true positives on average across samples from the Genome in a Bottle (GiaB) project (Zook et al., 2019) and the Human Genome Structural Variation (HGSV) consortium (Chaisson et al., 2019). Code and models for Samplot-ML are open source and freely available at github.com/mchowdh200/samplot-ml.

## 2. Related Work

### SV-Plaudit

As described above, Samplot makes it easy to be able to verify whether or not a putative SV is a True positive. SV-Plaudit (Belyeu et al., 2018) is a framework built on top of Samplot and Amazon Web Services to enable manual curation of SVs using a simple web interface. SV-plaudit can output a score for each reported SV based on how many annotators labelled a region as a true positive or false positive.

**Figure 1.**
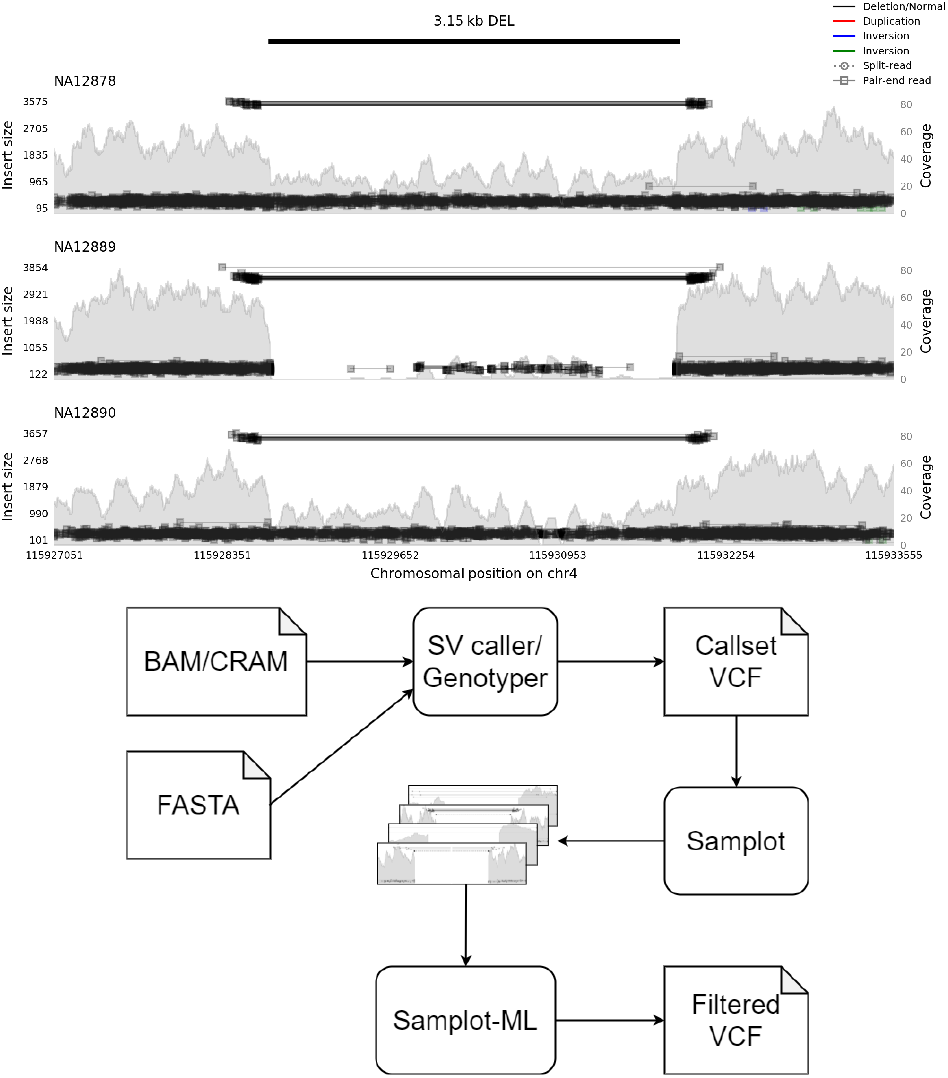
Top: Samplot images depicting SV calls. Low coverage, and discordant read pairs (black lines), are evidence of a deletion. Bottom: Typical workflow for Samplot-ML.

### Duphold

Many SV callers make use of discordant and split reads but do not incorporate depth of coverage. Duphold (Pedersen & Quinlan, 2019) is a heuristic-based method for filtering false-positive duplications and deletions. For each input region, Duphold computes a variety of metrics, including DHFFC (Duphold Flanking Fold Change), which, as the name suggests, computes the fold change in coverage between the reported region and the flanking regions.

### SV^2^

SV^2^ (Antaki et al., 2018) is a support vector machine trained on SV data from the 1000 Genomes Project (1000 Genomes Project Consortium et al., 2015) that genotypes duplications and deletions. SV^2^ extracts various features from each region such as depth of coverage, number of discordant/split reads, and the heterozygous SNV ratio.

## 3. Methods

### 3.1. Training Set

Our model was trained on data from 1,000 Genomes Project (1kg)(1000 Genomes Project Consortium et al., 2015), including the phase three SV call set and the newer high coverage alignments. We excluded individuals present in or related to individuals in our test sets (NA12878, NA12891, NA12892, HG00512, HG00513, HG00731, HG00732, NA19238, NA19239)

#### True Positive Regions

Heterozygous and homozygous deletions were sampled from the GRCh38 liftover of the phase 3 integrated SV map (Sudmant et al., 2015). Although this set contains high confidence SV calls, there were still a few false positives. To minimize the possibility of sampling a false positive, we filter this set using Dupholds DHFFC metric (ie. remove regions with DHFFC > 0.7). After filtering, we sampled 150,000 heterozygous deletions and 50,000 homozygous deletions.

#### True Negative Regions

Care must be taken to sample “true negatives” properly. Before choosing a negative set, we must consider the use case of our model. In practice, our model will remove false positives from the output set of an SV caller or genotyper. That means that our model will encounter two different classes of regions: those containing real SVs and edge cases that confused the SV caller’s filters. While we could have sampled regions from homozygous reference samples in the 1kg calls (i.e., samples without deletions) to get “true negatives”, these regions would have had very few discordant alignments and level depths of coverage. Crucially, they would look nothing like the regions that we would want our model to filter.

We took a more principled approach to pick true negatives. Many SV callers have the option to provide a set of “exclude regions”, which prevents the caller from considering potentially problematic regions of the genome (Li, 2014). Since these regions are enriched for false positives, we used these regions’ calls as our true negatives. To get variants in these regions, we recalled SVs on the 1kg high coverage alignments using Lumpy(Layer et al., 2014) with SV-Typer(Chiang et al., 2015). We then selected areas in the resultant calls that intersected problematic regions. To ensure that no true positives were selected, we filtered out regions with a DHFFC ≤ 0.7. Finally, to construct our set of true negatives, we took roughly 35,000 “exclude regions” and 15,000 homozygous reference regions from the 1kg call SV call set.

### 3.2. Test Sets

For testing, we used alignments from four individuals from Genome in a Bottle (HG002) and the Human Genome Structural Variation consortium (HG00514, HG00733, NA19240). For HG002, we obtained the alignements from (Pedersen & Quinlan, 2019) and used the GiaB v0.6 gold standard vcf (Zook et al., 2019) as our truth set. For HG00514, HG00733, NA19240, we obtained alignments from ftp://ftp.1000genomes.ebi.ac.uk/vol1/ftp/data_collections/hgsv_sv_discovery/data/ and the truth set vcf from ftp://ftp.ncbi.nlm.nih.gov/pub/dbVar/data/Homo_sapiens/by_study/genotype/nstd152.

### 3.3. Image generation

To generate images in both the training and test sets, we used the following samplot command: `samplot.py -c $chrom -s $start -e $end --min_mqual 10 -t DEL -b $bam -o $out_file -r $fasta` Where chrom, start, end are the genomic region, bam is the alignment file, and fasta is the reference genome file. Additionally, for SVs with length > 5000 bases, we added `--zoom 1000` which only shows 1000 bp centered around each breakpoint. After an image is generated, we crop out the plot text and axes using imagemagik (The ImageMagick Development Team). Finally, before input into Samplot-ML, the vertical and horizontal dimensions are reduced by a factor of eight.

### 3.4. Model

Samplot-ML is a resnet (He et al., 2015) like model that takes a Samplot image of a putative deletion SV as input and predicts whether it is homozygous reference, heterozygous, or homozygous alternate. For model details, see Figure 2.

**Figure 2.**
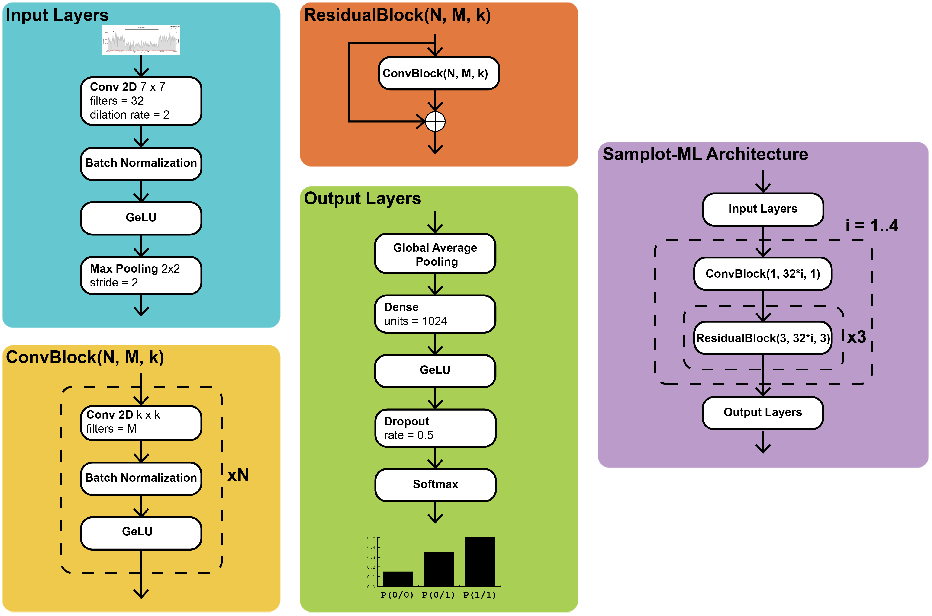
Samplot-ML model architecture. GeLU refers to the Gaussian error linear unit (Hendrycks & Gimpel, 2018).

### 3.5. Training Procedure

From our training set, we held out regions from chromosomes 1, 2, and 3 to use as a validation set. To train our model, we used stochastic gradient descent with warm restarts (SGDR) (Loshchilov & Hutter, 2017). The initial learning rate was 0.2 and decayed with a cosine annealing schedule. The initial restart period was set to two epochs and doubled after each restart. We trained for 50 epochs, and kept the model with the best validation loss after training was complete.

### 3.6. Testing Procedure

To evaluate the efficacy of Samplot-ML we called SVs and genotyped deletions using both Lumpy/SVTyper and Manta (Chen et al., 2016) on each of our test samples. We then filtered both Lumpy and Manta callsets with Duphold (rejecting calls with DHFFC ≤ 0.7), SV^2^, and Samplot-ML. To compare the filtered call sets, with their respective gold standards we used Truvari (tru, 2020), which compares regions in VCFs based on percent overlap as well as breakpoint accuracy. We used the following truvari command: `truvari -b $truth_set -c $filtered_call_set -o $out_dir --sizemax 1000000 --sizemin 300 --sizefilt 270 --pctovl 0.6 --refdist 20`.

## 4. Results

Using Samplot-ML, we were able to reduce false positives by 47% while preserving 97% of true positives on average across all test samples, consistently matching or beating Duphold and SV^2^ in F1 scores. For more detailed comparisons see Table 1 and Figure 3

**Figure 3.**
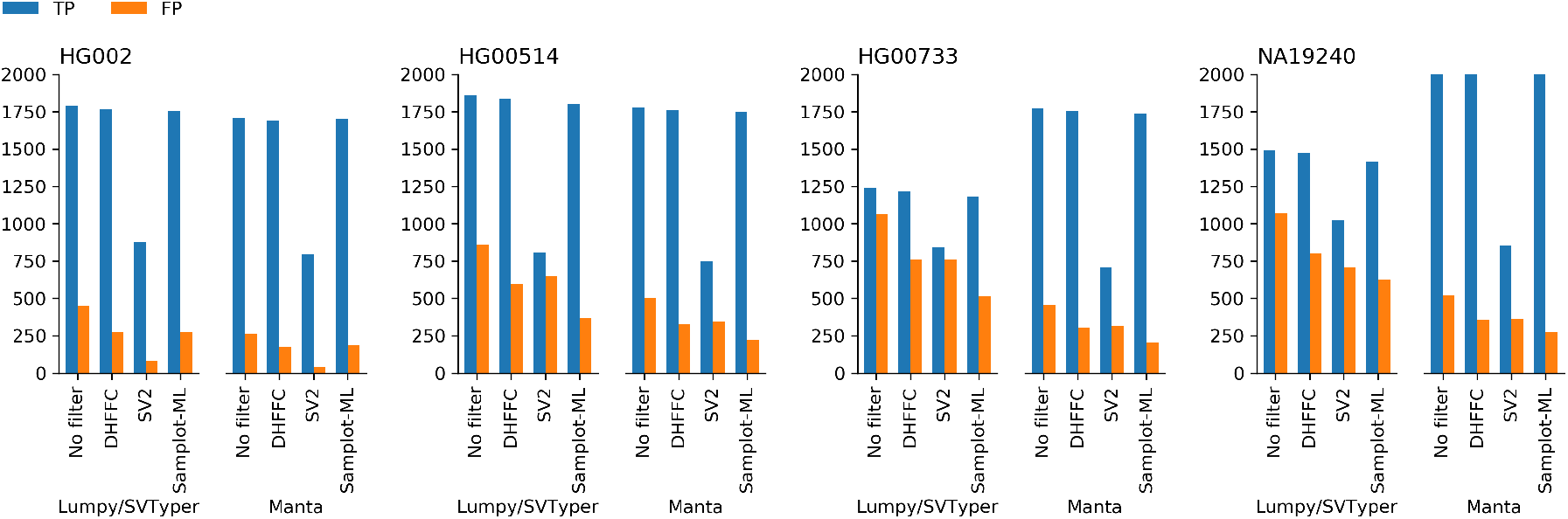
True positive and false postive comparisons between all methods.

**Table 1.** Complete Samplot-ML comparison statistics.

| Sample | Caller | Filter | TP | FP | FN | Precision | Recall | F1 |
| --- | --- | --- | --- | --- | --- | --- | --- | --- |
| HG002 | Lumpy/SVType | None | 1787 | 452 | 893 | 0.798 | 0.667 | 0.72 |
|  |  | DHFFC | 1764 | 276 | 916 | 0.865 | 0.658 | 0.747 |
|  |  | SV2 | 880 | 83 | 4012 | 0.914 | 0.180 | 0.301 |
|  |  | Samplot-ML | 1758 | 273 | 922 | 0.866 | 0.656 | 0.746 |
|  | Manta | None | 1708 | 265 | 972 | 0.866 | 0.637 | 0.734 |
|  |  | DHFFC | 1687 | 175 | 993 | 0.906 | 0.629 | 0.743 |
|  |  | SV2 | 799 | 41 | 4093 | 0.951 | 0.163 | 0.279 |
|  |  | Samplot-ML | 1699 | 187 | 981 | 0.901 | 0.634 | 0.744 |
| HG00514 | Lumpy/SVType | None | 1860 | 860 | 858 | 0.684 | 0.684 | 0.684 |
|  |  | DHFFC | 1837 | 596 | 881 | 0.755 | 0.676 | 0.713 |
|  |  | SV2 | 811 | 652 | 1907 | 0.554 | 0.298 | 0.388 |
|  |  | Samplot-ML | 1803 | 372 | 915 | 0.829 | 0.663 | 0.737 |
|  | Manta | None | 1779 | 502 | 939 | 0.780 | 0.654 | 0.712 |
|  |  | DHFFC | 1759 | 328 | 959 | 0.843 | 0.647 | 0.731 |
|  |  | SV2 | 748 | 345 | 1970 | 0.684 | 0.275 | 0.393 |
|  |  | Samplot-ML | 1751 | 221 | 967 | 0.888 | 0.644 | 0.747 |
| HG00733 | Lumpy/SVType | None | 1236 | 1066 | 1505 | 0.537 | 0.451 | 0.490 |
|  |  | DHFFC | 1216 | 760 | 1525 | 0.615 | 0.443 | 0.520 |
|  |  | SV2 | 842 | 760 | 1899 | 0.526 | 0.307 | 0.388 |
|  |  | Samplot-ML | 1181 | 517 | 1560 | 0.696 | 0.431 | 0.532 |
|  | Manta | None | 1774 | 455 | 967 | 0.796 | 0.647 | 0.714 |
|  |  | DHFFC | 1753 | 306 | 988 | 0.851 | 0.640 | 0.730 |
|  |  | SV2 | 707 | 317 | 2034 | 0.690 | 0.258 | 0.376 |
|  |  | Samplot-ML | 1736 | 204 | 1005 | 0.895 | 0.633 | 0.741 |
| NA19240 | Lumpy/SVType | None | 1494 | 1070 | 1711 | 0.583 | 0.566 | 0.518 |
|  |  | DHFFC | 1470 | 801 | 1735 | 0.647 | 0.459 | 0.537 |
|  |  | SV2 | 1025 | 708 | 2180 | 0.591 | 0.320 | 0.415 |
|  |  | Samplot-ML | 1414 | 628 | 1791 | 0.692 | 0.441 | 0.539 |
|  | Manta | None | 2067 | 520 | 1138 | 0.799 | 0.645 | 0.714 |
|  |  | DHFFC | 2054 | 359 | 1151 | 0.851 | 0.641 | 0.731 |
|  |  | SV2 | 855 | 362 | 2350 | 0.703 | 0.267 | 0.387 |
|  |  | Samplot-ML | 2019 | 272 | 1186 | 0.881 | 0.630 | 0.735 |

## 5. Conclusion

We present Samplot-ML, a convolutional neural network model that filters out potential false-positive deletions in SV call sets. Samplot-ML outperformed Duphold and SV^2^.

In many ways training a CNN to discriminate between different classes of images is relatively straightforward, given the current state of the art. The real challenge is in selecting positive and negative training examples that accurately reflect what real-world users will ask the model to classify. Data repositories like the 1,000 Genomes Project and ENCODE provide VCF or BED files that describe where genomic features occur (e.g., structural variants, regulatory elements, etc.). In the context of our classification task, these are good positive training examples. But, to accurately distinguish between a true positive and a false positive, we must also sample good negative examples.

In genome feature detection broadly, and SV detection specifically, negatives far outnumber positives. To achieve maximum classification performance, collecting negative training examples must be given as much consideration as any other aspect of the machine learning architecture. Just as it is highly unlikely that any genomic detection algorithm would return a random genomic region as a putative event, we cannot expect that randomly sampled areas of the genome that do not overlap true positives will be good negative examples. Special care must be taken to sample from regions enriched with edge cases that pass detection filters but do not contain true positives. By incorporating putative false positive areas of the genome, we were able to improve the performance of Samplot-ML immensely because these regions strongly resembled the types of false positives that were being made by SV callers.

## Notes

### Competing Interest Statement

The authors have declared no competing interest.

